# Impaired Sensory Gating During Standing Balance in Parkinson’s Disease

**DOI:** 10.1101/2025.08.21.668914

**Authors:** Ashwini Sansare, Hossein Soroushi, Taylor M Gauss, Jan M Hondzinski, Deanna Kennedy, Yuming Lei

## Abstract

While motor deficits in Parkinson’s disease (PD) are well-studied, the role of somatosensory processing in postural instability remains unclear. It is unknown whether sensory gating, a mechanism for filtering irrelevant sensory input, is impaired during standing balance in individuals with PD. To address this, we investigated cortical sensory processing in individuals with PD, age-matched older adults (OA), and young adults (YA) as they performed four balance tasks of increasing difficulty. We measured postural sway using a force platform and recorded somatosensory-evoked potentials (SEPs) from the primary somatosensory cortex (S1) following tibial nerve stimulation. Our results showed a clear dissociation between behavior and neurophysiology. Although postural sway was comparable between the PD and OA groups, only the OA and YA groups showed intact sensory gating, with SEP amplitudes decreasing as the balance challenge increased. In contrast, participants with PD demonstrated consistently elevated SEP amplitudes across all conditions. This study provides the first direct evidence of impaired sensory gating during standing balance in PD. These findings indicate a fundamental deficit in the cortical processing of sensory information essential for postural control. Consequently, they underscore the critical need for therapeutic interventions that target sensory integration deficits, not just motor symptoms.

**Key points:**

- Healthy young and older adults demonstrate intact sensory gating during standing balance, with somatosensory-evoked potential (SEP) amplitudes decreasing as postural difficulty increases.
- Individuals with Parkinson’s disease (PD) show impaired sensory gating, with elevated SEP amplitudes that are not appropriately modulated by increasing postural demands.
- Despite comparable postural sway to healthy older adults, the PD group exhibited fundamentally different neurophysiological responses to balance challenges.
- This dissociation between motor performance and neurophysiology indicates a primary deficit in cortical sensory processing in PD.
- Impaired sensory gating may reflect a key, independent contributor to postural instability in PD, highlighting the need to target sensory deficits in treatment.

## Introduction

Almost 60% of individuals with Parkinson’s Disease (PD) fall at least once every 3-6 months, with 40% of them falling recurrently (1). The devastating consequences include injury, reduced activity levels and quality of life (2), and an increased fear of falling (3,4). Given that the prevalence of PD is expected to double by 2030 (5), addressing the root causes of postural instability is a critical priority. While research has traditionally emphasized motor deficits, the contribution of sensory processing to balance impairment is less understood. Therefore, in this study we aim to determine how PD alters the sensory mechanisms involved in maintaining postural control.

Postural instability in PD arises from a complex interplay of motor and sensory deficits. While the motor component, driven by dysfunction in regions like the primary motor area (M1) and the basal ganglia (6–9), has been extensively studied, the contribution of sensory dysfunction, increasingly recognized as a critical factor for control, has not. Central to this is the primary somatosensory cortex (S1), which performs a critical filtering function known as sensory gating. This mechanism dynamically suppresses sensory inputs irrelevant to stability, ensuring that only the most pertinent information guides postural corrections (10–12). The importance of this is underscored by the wealth of evidence demonstrating sensory impairments in PD, including reduced tactile and spatial acuity, altered pain and temperature sensitivity, and significant proprioceptive deficits (13,14). A leading hypothesis for these deficits is that dopaminergic dysfunction in the basal ganglia disrupts key sensory processing hubs, including the thalamus and S1, a theory supported by evidence of significant thalamic neuronal loss in PD (15,16). However, the exact neurophysiological mechanisms through which these somatosensory deficits impair postural control remain unknown.

To objectively measure the transmission and filtering of sensory information, we utilized somatosensory-evoked potential (SEP). Distinct SEP components mark the arrival of sensory signals at key structures like the thalamus and S1 (17–20). Crucially, SEP amplitude is not static but is dynamically modulated during movement through sensory gating. This filtering is indexed by an attenuation of the SEP amplitude, with a greater reduction signifying more effective gating (21). For instance, our previous studies showed that healthy adults exhibit adaptive gating, with SEP amplitudes decreasing as upper-limb task difficulty increases (22,23).

This adaptive mechanism of sensory gating appears to be compromised in PD. Indeed, correlations between altered SEPs and poor upper limb function in PD suggest a fundamental deficit in sensorimotor processing (24–26). However, whether this gating impairment extends to the critical task of balance control remains unknown. Therefore, we use the present study to determine whether these gating deficits affect balance control in PD. We investigated sensory gating during standing tasks of increasing difficulty in individuals with PD and healthy control groups, hypothesizing that the PD group would exhibit impaired sensory gating, characterized by an inability to modulate SEP amplitude effectively in response to increasing postural demands.

## Materials and Methods

### Participants

Three groups of sixteen individuals participated: individuals with PD (Hoehn & Yahr Stage II; mean age = 69, SD = 7, 6 females), age-(±6 months) and sex-matched healthy older adults (OA; mean age = 69, SD = 6, 6 females), and healthy young adults (YA; mean age = 25, SD = 3, 10 females). The local Institutional Review Board approved the experimental protocol, and all participants provided written informed consent. The study was performed in accordance with the Declaration of Helsinki. Inclusion criteria for the PD group included a prior medical diagnosis, age 40-80 years, and the ability to stand unassisted for at least 30 s. Control participants had no known medical or neurological conditions. Participants were excluded for the presence of metal implants affecting recordings, any neurological disease other than PD (for the PD group only), recent lower limb injuries or surgeries (within 2 years), or unstable medications affecting balance. All participants with PD were tested while on their regular medication schedule.

### Balance Assessment

Postural sway was recorded using an Accusway force platform (Advanced Mechanical Technology Inc., Watertown, MA). A medium-density foam pad was used for unstable surface conditions. Participants were instructed to stand as still as possible with their feet positioned 10 cm apart. They completed trials under four conditions, presented in a randomized order: (1) eyes open on a firm surface (EO), (2) eyes closed on a firm surface (EC), (3) eyes open on a foam surface (EOF), and (4) eyes closed on a foam surface (ECF) (Figure 1). To mitigate potential electrical interference between the force platform and the EEG system, each 30-second balance trial was immediately followed by a 30-second period of quiet standing for EEG recording with the force platform unplugged. Participants completed three such trial pairs for each of the four conditions. Center of pressure (COP) data were sampled at 100 Hz and then low-pass filtered using a 9th order Butterworth filter with a 20 Hz cutoff. The primary measure for postural sway was the 95% confidence ellipse area (95% COP area), which encompasses 95% of the samples drawn from the underlying distribution. Secondary measures included the root mean square (RMS) of COP displacement in the anterior-posterior (A-P) and medial-lateral (M-L) directions, mean COP velocity, and COP frequency dispersion (27–31).

**Figure 1:**
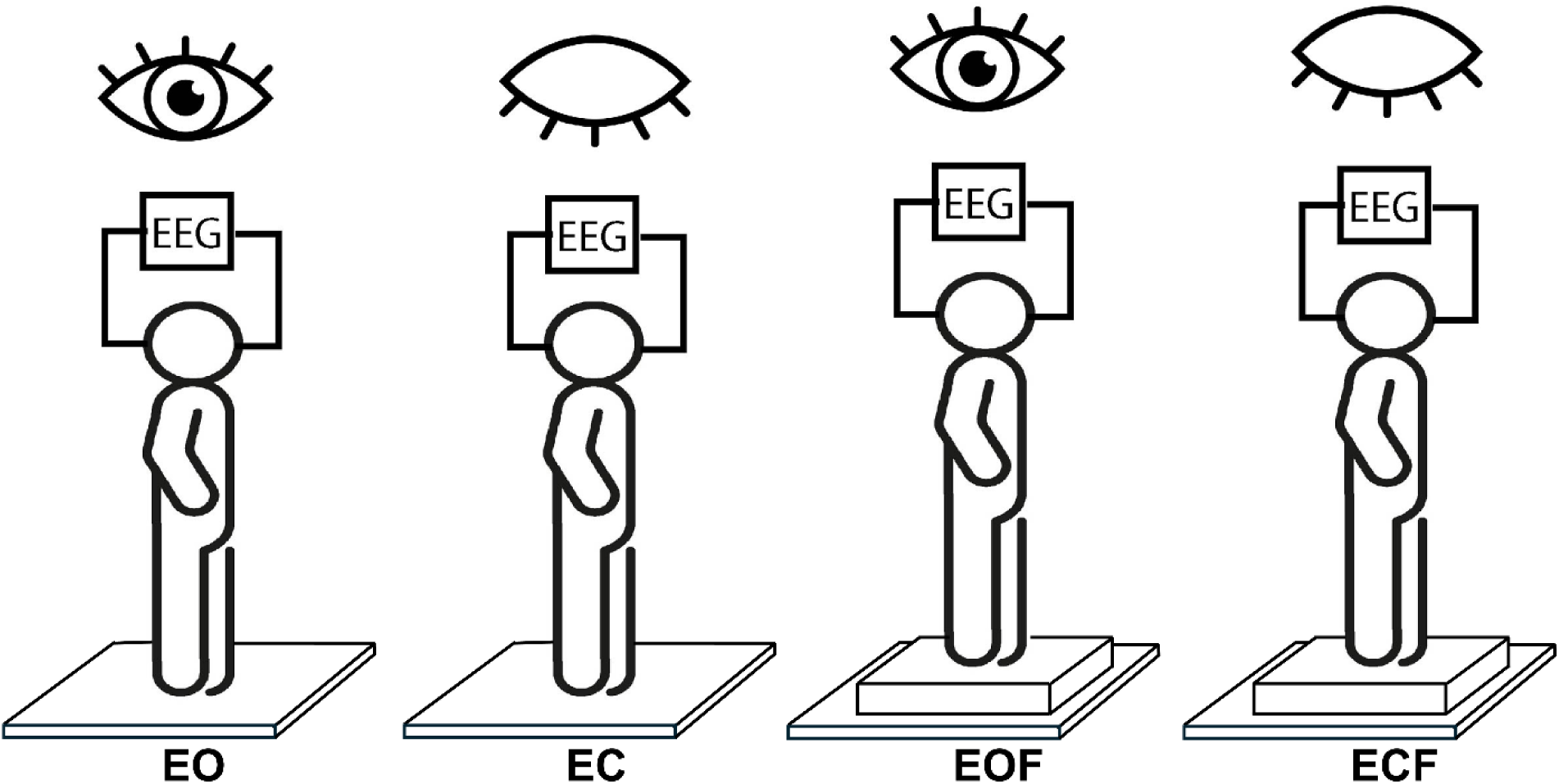
Participants performed four different balance conditions; eyes open (EO), eyes closed (EC), eyes open on foam surface (EOF), eyes closed on foam surface (ECF) while standing on a force plate. SEPs were measured using electroencephalography (EEG) during these four conditions.

### Somatosensory Evoked Potentials (SEPs)

SEPs were recorded from the left primary somatosensory cortex (S1) using a wearable 4-channel EEG system (NeXus-10 MKII, Mind Media, Netherlands). To elicit the SEPs, peripheral electrical stimulation (10 ms pulse duration) was applied to the tibial nerve just posterior to the medial malleolus at a frequency of 5 Hz. For each condition, participants received up to 450 stimulation pulses. Stimulation intensity was set to three times each individual’s sensory threshold while remaining below their motor threshold, ensuring a perceptible stimulus without inducing muscle activation. Sensory threshold was defined as the minimum level of stimulation required for an individual to detect a tingling sensation at the stimulation site. The stimulation intensity was increased in increments of 0.1 mA, starting at zero, until the participant reported feeling the stimulation. To verify this threshold, the intensity was decreased until the participant could no longer feel the stimulation. This procedure was repeated three times for each participant, and the sensory threshold for that site was defined as the lowest value over the three repetitions.

EEG electrodes were positioned according to a modified 10–20 system: 5 cm lateral and 5 cm anterior to the vertex to target the frontal component, and 2 cm posterior to the vertex for the parietal component, corresponding to regions just anterior and posterior to C3 in the standard layout. The ground electrode was placed on the forehead, and reference electrodes were positioned on the bilateral mastoid processes—an arrangement previously validated for SEP recording in our previous paradigms (22,23).

All preprocessing was performed in EEGLAB (32) with custom MATLAB scripts (The MathWorks, Natick, MA). SEP preprocessing followed published procedures (33). EEG data were sampled at 2,048 Hz. Stimulation artifacts occurring 4–8 ms post-stimulus were removed using piecewise cubic Hermite interpolation (PCHIP). Signals were then band-pass filtered (30–200 Hz) with a zero-phase, 4th-order Butterworth filter applied forward and reverse to avoid phase distortion and to attenuate slow drifts; 60-Hz line noise was removed with a 5-Hz-bandwidth notch filter. To mitigate variability from electrode impedance and reference instability, we applied a weighted subtraction procedure: channel-specific weights and y-intercepts were estimated to minimize the mean over the pre-stimulus baseline, yielding baseline-centered traces. Data were visually inspected and segments with eye movements or high-frequency muscle activity were rejected, resulting in an average of ∼300 valid epochs per participant. Epochs spanned −50 to 100 ms around stimulus onset, with baseline correction using the −50 to 0 ms interval (32). To quantify S1 activation, we measured the P38 SEP component, selected for its well-established generator in the contralateral primary somatosensory cortex (20,34). For each participant and condition, P38 amplitude was computed as the average peak-to-peak response across all valid trials, providing a robust index of cortical sensory processing.

## Statistical analysis

All statistical analyses were performed using IBM SPSS Statistics. To assess the effects of group and condition on our primary outcome measures (95% COP Area and P38 SEP Amplitude), we conducted a series of 3 (Group: PD, OA, YA) × 4 (Condition: EO, EC, EOF, ECF) mixed-model analyses of variance (ANOVAs), with Group as the between-subjects factor and Condition as the within-subjects factor. Prior to analysis, assumptions of normality and homoscedasticity were confirmed using Shapiro-Wilk and Levene’s tests, respectively. Mauchly’s test was used to assess the assumption of sphericity; in cases where this assumption was violated, the Greenhouse-Geisser correction was applied. Significant interactions were followed up with pairwise post-hoc comparisons using a Bonferroni correction for multiple comparisons. The significance level for all tests was set at p < 0.05.

## Results

From the initial sample of 48 participants, data from five individuals were excluded from the final SEP analysis due to poor signal quality or unreliable determination of the P38 component (2 PD, 1 OA, 1 YA). Additionally, one OA participant was identified as a statistical outlier in the balance analysis, with postural sway values more than double the group average. The final data sets used for the primary analyses therefore included 14 PD, 14 OA, and 15 YA participants. For the sake of complete transparency, all analyses were repeated including the outlier, and these results are presented in Supplementary File 1.

Table 1 shows the descriptive statistics for 95% COP area and SEP.

**Table 1:** Mean and Standard deviations for balance and SEP, split by group and condition.

|  | Condition | Group | n | Mean | Std Dev |
| --- | --- | --- | --- | --- | --- |
| 95%COParea (sq.cm) | EO | PD | 16 | 3.01 | 5.17 |
|  |  | OA | 16 | 1.71 | 1.04 |
|  |  | YA | 16 | 1.51 | 1.41 |
|  | EC | PD | 16 | 2.81 | 1.75 |
|  |  | OA | 16 | 2.43 | 1.27 |
|  |  | YA | 16 | 1.97 | 2.30 |
|  | EOF | PD | 16 | 10.81 | 10.54 |
|  |  | OA | 16 | 7.35 | 2.99 |
|  |  | YA | 16 | 4.56 | 1.75 |
|  | ECF | PD | 16 | 18.50 | 10.65 |
|  |  | OA | 16 | 20.85 | 9.32 |
|  |  | YA | 16 | 9.19 | 2.55 |
| SEP | EO | PD | 14 | 2.95 | 1.27 |
|  |  | OA | 13 | 1.98 | 0.54 |
|  |  | YA | 15 | 1.90 | 0.56 |
|  | EC | PD | 14 | 2.78 | 1.19 |
|  |  | OA | 13 | 1.67 | 0.49 |
|  |  | YA | 15 | 1.49 | 0.41 |
|  | EOF | PD | 14 | 2.60 | 1.22 |
|  |  | OA | 13 | 1.44 | 0.37 |
|  |  | YA | 15 | 1.58 | 0.44 |
|  | ECF | PD | 14 | 3.35 | 1.13 |
|  |  | OA | 13 | 1.29 | 0.38 |
|  |  | YA | 15 | 1.30 | 0.32 |

### Balance Assessment

The 3 (Group) × 4 (Condition) mixed-model ANOVA on 95% COP area revealed a significant Group × Condition interaction (F= 5.205, p = 0.003, η2 = 0.188), indicating that the effect of the balance conditions on postural sway differed among the three groups. Figure 2 shows results of the pairwise post hoc comparisons using Bonferroni corrections. During the simpler firm-surface conditions (EO and EC), there were no significant differences in postural sway among the three groups. Group differences only emerged as the task became more challenging on the unstable foam surface. In the eyes-open on foam (EOF) condition, the PD group showed a significantly higher postural sway compared to the YA group (p = 0.025). This effect was magnified in the most difficult eyes-closed on foam (ECF) condition, where both the PD group (p = 0.008) and the OA group (p < 0.001) swayed significantly more than the YA group. Notably, no significant differences in postural sway were found between the PD and OA groups in any of the four conditions. This overall pattern was consistent across our secondary sway measures (e.g., COP velocity, RMS), with full details provided in Supplementary File 2.

**Figure 2.**
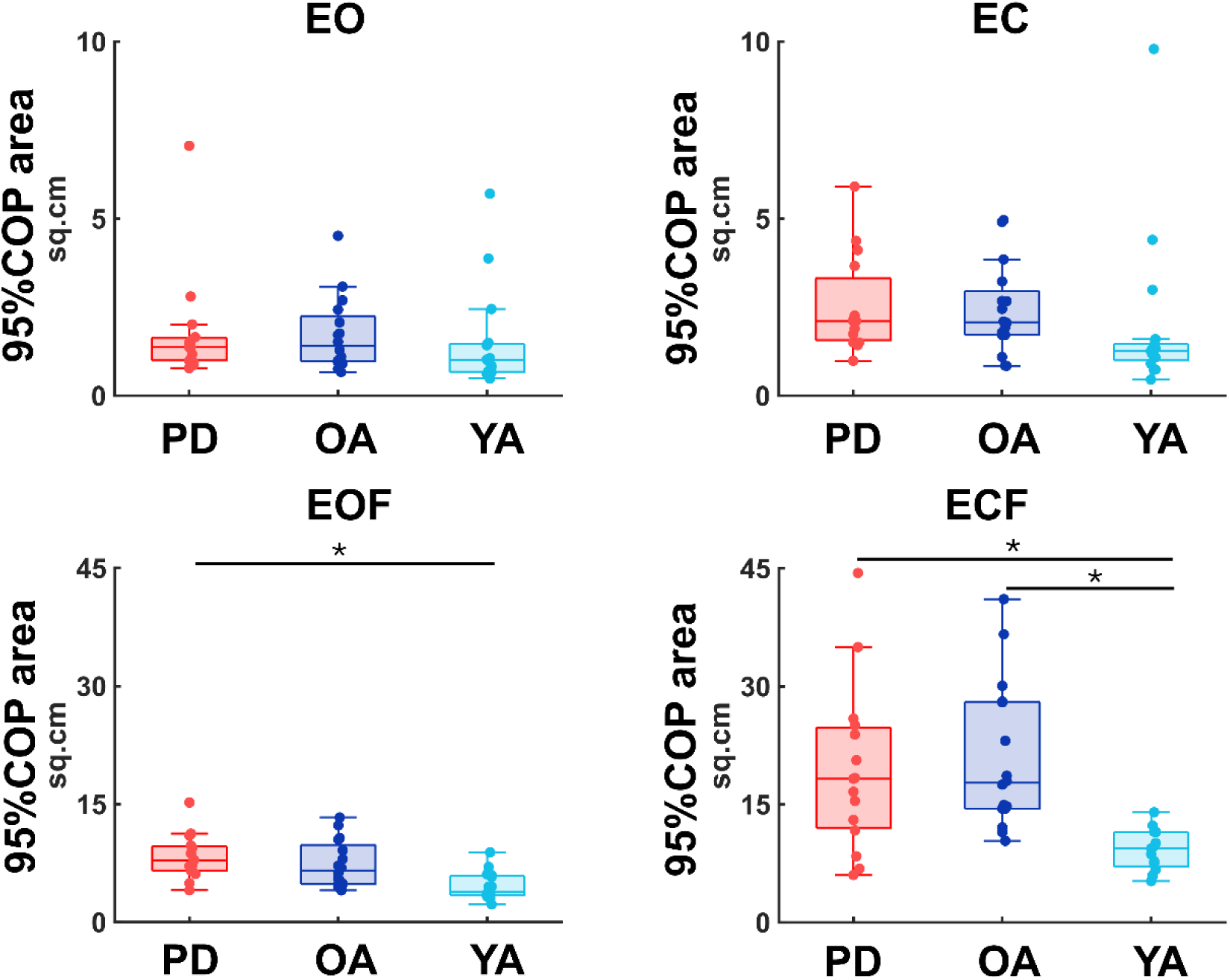
Box and whisker plots, with scattered dots indicating each subject showing differences in the 95% COP area between the three groups, PD (red), OA (dark blue) and YA (light blue) groups across the four balance conditions, eyes open (EO), eyes closed (EC), eyes open on foam (EOF), and eyes closed on foam (ECF) shown in each panel. Asterisks above horizontal segments represent significant differences between means (p < 0.05).

Figure 3 shows results of the Bonferroni post-hoc comparisons for postural sway across conditions within each group. While postural sway during ECF significantly exceeded EC (p = 0.004) and EO (p = 0.012) for the YA group, it exceeded EOF (p < 0.001), EC (p < 0.001), and EO (p < 0.001) for the OA group and EOF (p = 0.031), EC (p < 0.001), and EO (p < 0.001) for the PD group. Additionally, although postural sway during EOF significantly exceeded EO (p = 0.009) for the YA group, it exceeded EC (p = 0.007) and EO (p < 0.001) for the OA group and EC (p < 0.001) and EO (p < 0.001) for the PD group. These results confirm that while all groups found the tasks progressively more difficult, the older and PD groups were more affected by the combined loss of reliable visual and somatosensory information.

**Figure 3.**
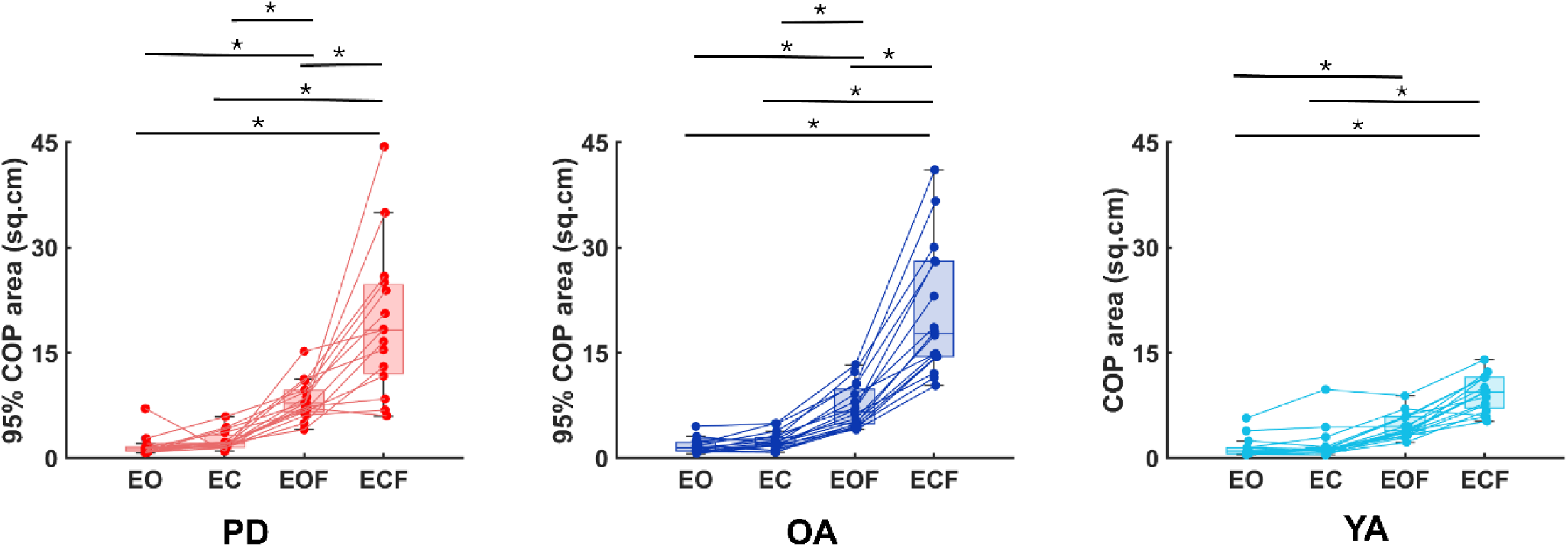
Box and whisker plots, with scattered dots indicating each subject showing the trend in the 95% COP area as the balance conditions become increasingly challenging from eyes open (EO), eyes closed (EC), eyes open on foam (EOF) to eyes closed on foam (ECF) for the three group, PD (red), OA (dark blue) and YA (light blue). Asterisks above horizontal segments represent significant differences between means (p < 0.05).

### SEPs

The 3 (Group) × 4 (Condition) mixed-model ANOVA on SEP amplitude revealed a significant Group × Condition interaction (F= 9.301, p < 0.001, η2 = 0.323). This indicates that the groups’ cortical sensory responses were modulated differently by the balance conditions. Post-hoc comparisons revealed a clear and consistent main effect of group (Figure 4). The between group comparisons for PD, OA and YA for each condition showed that SEP amplitude for the PD group significantly exceeded the OA group (p = 0.018 for EO, p = 0.002 for EC, p = 0.001 for EOF, p < 0.001 for ECF) and YA group (p = 0.007 for EO, p < 0.001 for EC, p = 0.003 for EOF, p < 0.001 for ECF). This finding points to a persistent, over-activation of the somatosensory cortex in the PD group that is present regardless of immediate postural demands.

**Figure 4.**
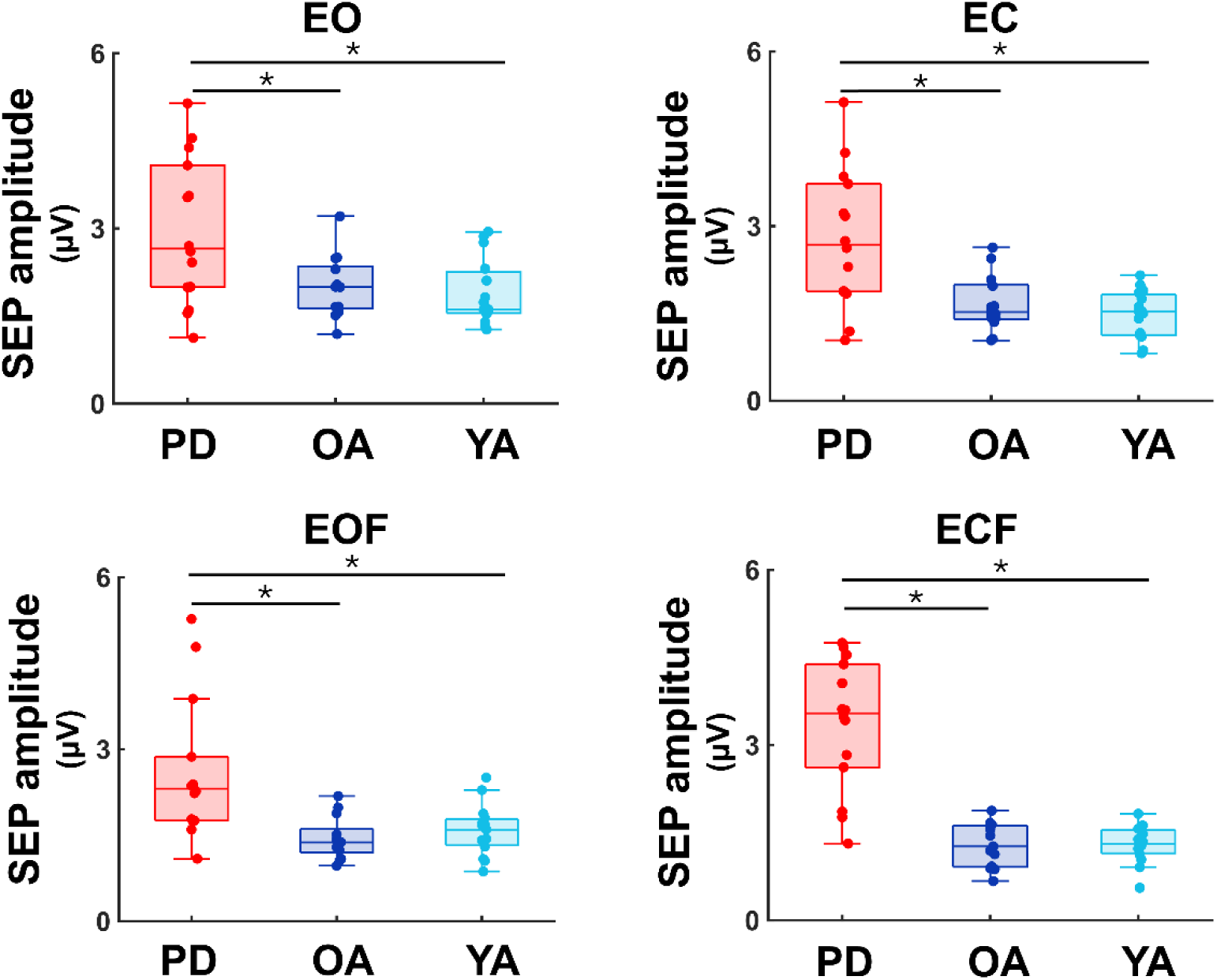
Box and whisker plots, with scattered dots indicating each subject showing differences in the SEP amplitude between the three groups, PD (red), OA (dark blue) and YA (light blue) groups across the four balance conditions, eyes open (EO), eyes closed (EC), eyes open on foam (EOF) and eyes closed on foam (ECF) shown in each panel. Asterisks above horizontal segments represent significant differences between means (p < 0.05).

The analysis of how SEP amplitudes changed across conditions within each group revealed fundamentally different patterns of sensory modulation (Figure 5). Both the YA and OA groups demonstrated adaptive sensory gating. Their SEP amplitudes were highest in the simplest condition (EO) and were significantly attenuated as the balance tasks became more challenging (e.g., for OA, EO vs. ECF, p < 0.001). This reduction in amplitude reflects the successful filtering of the irrelevant electrical stimulus as the need to focus on postural sensory cues increased. In contrast, the PD group failed to show this adaptive modulation. Their SEP amplitudes were not suppressed with increasing task difficulty. Instead, the amplitude in the most difficult condition (ECF) was significantly higher than in the easier conditions (e.g., vs. EO, p < 0.001). These opposing patterns—attenuation in healthy controls versus amplification in the PD group under maximal sensory conflict—provide strong evidence for a dysfunctional sensory gating mechanism in PD.

**Figure 5.**
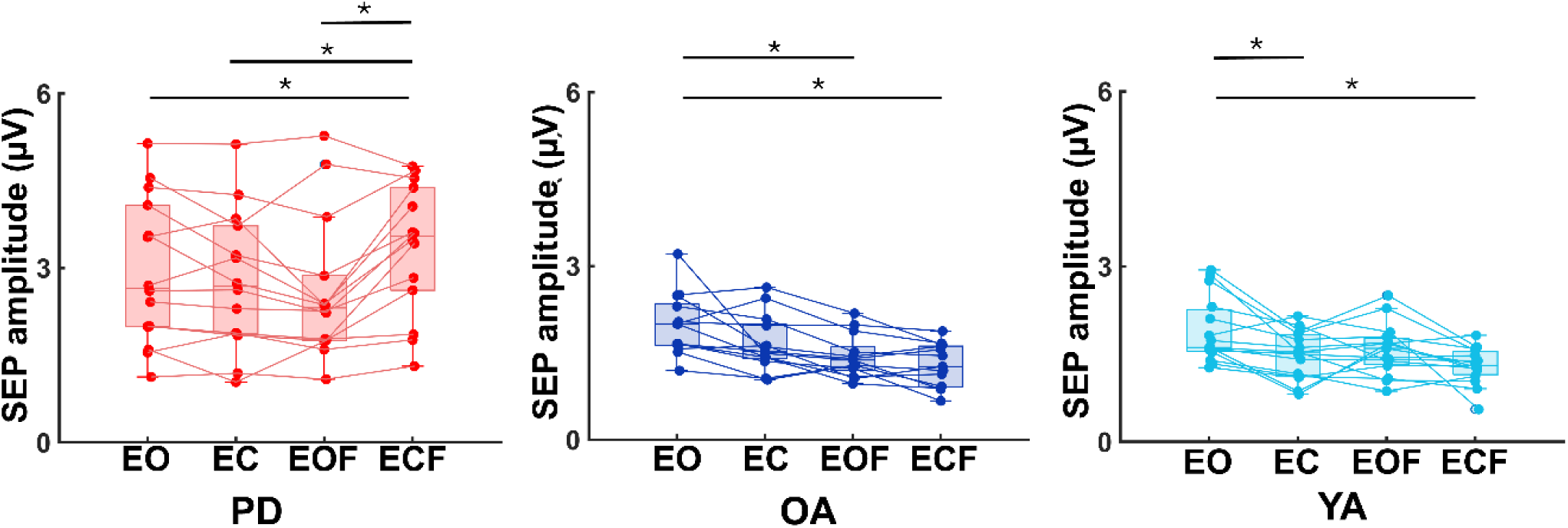
Box and whisker plots, with scattered dots indicating each subject showing the trend in the SEP amplitude as the balance conditions become increasingly challenging from eyes open (EO), eyes closed (EC), eyes open on foam (EOF) to eyes closed on foam (ECF) for the three group, PD (red), OA (dark blue) and YA (light blue). Asterisks above horizontal segments represent significant differences between means(p < 0.05).

## Discussion

We examined the gating of sensory input during increasingly difficult balance tasks in individuals with PD, age-and sex-matched healthy older adults, and healthy young adults. We found evidence for significantly worse postural sway in PD and OA groups compared to the YA group in the most challenging condition. We also found that as balance conditions got progressively more challenging, the SEP amplitude reduced in the YA and OA group, showing that the sensory cortex was able to gate the irrelevant sensory input delivered via the external electrical stimulation. Conversely, the greater increase in SEP amplitude on the most difficult condition and no significant difference in the SEP amplitude across the remaining conditions for the PD group indicated altered processing of the sensory input and less ability to gate irrelevant sensory input. Given the comparable postural sway between the OA and PD groups, the differences in the sensory gating highlight the fact that sensory processing impairments from the lower limb in our PD group did not need to impair motor performance in postural sway.

### Postural Sway in PD and OA Exceeded YA

Our results indicated that the OA group exhibited significantly greater postural sway than the YA group only during the most challenging balance condition. This finding aligns with previous research showing that postural sway increases with age (35,36), particularly under conditions without visual input and altered proprioceptive feedback through changes in the support surface (37,38). Balance abilities used for postural sway clearly decline with age without access to reliable sensory inputs from visual and proprioceptive systems. In contrast, the PD group’s postural instability was more pronounced, showing significantly greater sway than the YA group as soon as proprioceptive feedback was challenged on the foam surface, regardless of visual condition. This pattern is consistent with the known pathophysiology of PD, where basal ganglia degeneration impairs the ability to automatically regulate and scale postural responses (39–41).

Interestingly, we found no significant differences in postural sway between the PD and OA groups. We propose two potential explanations for this finding. First, the PD group comprised a high-functioning cohort, many of whom maintain functionally independence and engage in regular physical activity. Second, PD participants assessed on medication may reveal high function during motor performances not observed when medication wears off. Since most individuals with PD spend the majority of their day in the medicated state, this approach provided ecologically valid insights into their ordinary performance without imposing uncomfortable or distressing risks to participants’ safety. Therefore, our findings suggest that when optimally medicated, high-functioning individuals with PD can achieve a level of postural control that is comparable to their healthy, age-matched peers.

### YA and OA Show Adaptive, Age-Dependent Sensory Gating

YA and OA demonstrated effective sensory gating, a mechanism for filtering irrelevant sensory information that is critical for optimizing balance control. This process is indexed neurophysiologically by the attenuation of SEP amplitude with a greater reduction signifying more robust filtering of the external sensory stimulus as the demands of the postural task increase. Our results revealed distinct, age-dependent strategies for this gating. YA exhibited a gating strategy triggered primarily by the removal of vision; their SEP amplitudes decreased significantly once visual input was removed (EO > EC). In contrast, OA showed a more pronounced gating response that was sensitive to proprioceptive challenges; their SEP amplitudes decreased significantly when standing on foam (EO > EOF). Notice that this attenuation with the removal of a single sensory modality (EC or EOF) for either group did not correspond to behavioral declines in postural control, suggesting use of compensatory mechanisms by both groups. However, attenuation of proprioceptive feedback from the unstable foam surface and visual feedback simultaneously (ECF) provided the strongest driver for sensory gating, especially affecting the ability of the OA group to compensate and limit their postural sway. Researchers attribute the inability of older people to counteract reductions in multiple sensory inputs to increased attentional resources compared to their younger counterparts used for balance control (42), eliminating the use of more automated and successful control of posture (43). These findings of adaptive, age-specific gating strategies also provide a crucial neurophysiological baseline against which the deficits in PD can be understood.

The P38 SEP amplitude investigated here likely reflects the activation of S1 in response to sensory input from tibial nerve stimulation (20). As one of the earliest cortical areas to receive peripheral somatosensory input, S1 plays a critical role in initial sensory processing (44–47). The observed reduction in SEP amplitudes during the EOF and ECF conditions for healthy older adults indicates that, through altered proprioceptive input alone or in combination with removal of visual input—S1 selectively gated the incoming sensory signal. This likely reflects the brain’s prioritization of behaviorally relevant sensory inputs during postural control and suppression of irrelevant or redundant information to optimize balance performance.

### Impaired and Maladaptive Sensory Processing in PD

In contrast to the adaptive gating seen in controls, the PD group exhibited a fundamentally dysfunctional pattern of sensory processing. This was evident in two key findings. First, the PD group had significantly higher SEP amplitudes than both control groups across all conditions, even during simple standing. This tonic over-activation of the sensory cortex suggests two potential underlying issues: 1) a primary gating deficit, where the parkinsonian brain is simply unable to suppress irrelevant sensory input, or 2) a maladaptive compensatory strategy, where the brain tonically increases the’gain’ on all sensory information to overcome underlying uncertainty. This latter strategy creates a critical’ceiling effect,’ leaving little neural reserve to respond to more demanding tasks and offering a compelling explanation for why individuals with PD are so vulnerable in dynamic environments.

Second, within the PD group, SEP amplitude during the ECF condition was significantly higher than in the other three conditions. At the systems level, several mechanisms could contribute to this heightened SEP amplitude. One possibility is cortical disinhibition, whereby reduced GABA-A–mediated intracortical inhibition, which is well documented even in early PD, removes the normal inhibitory control that limits cortical excitability (48). In the context of balance, this disinhibition could allow excessive somatosensory input to reach higher-order processing areas, overwhelming integration centers and making it harder to selectively attend to the most relevant cues for postural stability. Another, not mutually exclusive, explanation is a compensatory gain increase in early sensory processing. The central nervous system, when faced with unreliable or challenging sensory inputs, upregulates the “gain” on afferent signals to maximize the extraction of usable information. During ECF, when both vision and reliable somatosensory cues from the surface are removed, this gain increase might help detect subtle postural sway or perturbations. However, the trade-off is that higher gain amplifies both signal and noise, which could introduce instability by triggering inappropriate or exaggerated postural responses (13). This finding suggests that when balance is maximally challenged in individuals with PD, as in the ECF condition, there may be a breakdown in normal sensory gating mechanisms. Specifically, the elevated SEP amplitude could reflect a compensatory response in which the brain allows a greater volume of afferent input to reach cortical processing areas, possibly in an effort to maintain postural stability under extreme sensory conflict. In healthy individuals, sensory gating serves to filter out irrelevant or redundant information; however, in PD, where baseline sensory integration and processing is already impaired (49–52), this mechanism may become dysregulated. The absence of sensory gating under high-challenge conditions may result in a form of cortical “flooding” with peripheral input, as the central nervous system attempts to compensate for the loss of stable sensory references. Together, cortical disinhibition, and compensatory gain increase may operate in parallel, producing both pathological and adaptive changes in sensory processing. This combination could explain why SEP amplitudes are heightened in PD during maximal sensory conflict, reflecting a nervous system that is both overresponsive and inefficient in translating sensory input into stable postural control strategies.

### Dissociation of Neurophysiology and Behavior

Interestingly, while there were no significant group differences in postural sway between individuals with PD and age-matched OA, there were marked differences in SEP between these groups. This dissociation suggests that although dopaminergic medication may help restore motor function, gait and stabilize gross postural control in PD (53,54), it may not fully address the underlying sensory processing abnormalities. All PD participants in the present study were tested in their clinically defined ON-medication state for safety and ecological validity, which likely optimized motor performance. However, the persistence of SEP differences despite dopaminergic treatment is consistent with prior reports showing mixed or minimal effects of medication on somatosensory evoked potentials and sensory gating (55,56). This suggests that central somatosensory integration deficits in PD may be at least partly independent of dopamine-mediated pathways and therefore remain impaired even when overt behavioral performance appears normalized. These findings align with previous work indicating that sensory dysfunction in PD, particularly in proprioceptive and somatosensory processing, may persist regardless of motor symptom severity or medication status (55,57). Thus, neurophysiological measures such as SEP may offer unique insights into cortical-level dysfunction not readily apparent through behavioral assessment alone. This underscores the importance of incorporating multimodal approaches, combining behavioral and neurophysiological metrics, into clinical evaluations and interventions to address the full spectrum of sensorimotor impairments in PD.

While this work provides valuable insight into sensory processing during upright stance in individuals with PD, it contains some limitations. First, our EEG setup and protocol indirectly captured cortical activity and not subcortical structures such as the thalamus which directly contribute to sensory gating. Future studies which incorporate high-density EEG montages with greater capabilities could offer a more comprehensive understanding of sensory gating in subcortical structures during balance control in PD. These setups may improve spatial resolution and with a greater number of channels could enable more accurate source localization, thereby allowing simultaneous investigation of cortical and subcortical contributions to sensory gating mechanisms. Second, while SEP provided a valuable early measure of sensory processing, it reflected only one component of complex processes involved during sensorimotor control of balance. Other modalities, such as EEG-based connectivity, paired pulse transcranial magnetic stimulation approaches, and functional magnetic resonance imaging techniques may provide complementary insights. Lastly, most participants from the PD group were tested on medication. While ecologically valid, this may mask more pronounced deficits that could emerge without medication. Future studies could compare SEP responses off medication too.

## Conclusion

In conclusion, the results of this study revealed a stark dissociation between behavioral performance and cortical sensory processing in individuals with PD. While behavioral measures of postural sway were similar between individuals with PD and healthy older adults, their underlying neurophysiological responses were fundamentally different. Healthy controls successfully employed sensory gating, attenuating SEP amplitudes as balance tasks became more difficult. In contrast, the PD group failed to modulate their sensory response, instead showing a paradoxical *increase* in SEP amplitude under the most challenging condition. This demonstrates a clear disruption of normal gating mechanisms in PD. This dissociation provides compelling evidence that dopaminergic medication, while improving postural stability, may not fully restore underlying sensory integration processes. Our findings underscore a persistent sensory processing deficit in PD—a deficit not apparent from behavioral assessment alone but clearly revealed by neurophysiological measures like SEPs. These results contribute to a growing body of evidence on the importance of central sensory processing for balance control in neurodegenerative populations. Clinically, our findings suggest that therapeutic strategies for PD must expand beyond the motor domain to address these underlying sensory integration deficits, potentially through sensorimotor training or neurofeedback. Future research is needed to explore these interventions and to track how sensory gating dynamics evolve across disease stages and medication states.

