## Supplementary File 1 for "Impaired Sensory Gating During Standing Balance in Parkinson’s Disease"

Results from the 3 (group) x 4 (condition) ANOVA that included the outlier from the OA group revealed a significant group by condition interaction for SEP amplitude (F= 9.561, p < 0.001, ***η***^2^ = 0.323) (the analysis without this participant included in the main manuscript shows F= 9.301, p < 0.001, ***η***^2^ = 0.323). Supplementary Figure 1 shows the pairwise post hoc comparisons using Bonferroni corrections that include the outlier.

The between-group comparisons between PD, OA and YA for each condition showed that SEP amplitude for the PD group was not significantly higher than the OA group for EO (p = 0.188) and for EOF (p = 0.057). The SEP amplitude for PD group was significantly higher than OA for EC (p = 0.029) and for ECF (p < 0.001) and YA group (p = 0.022 for EO, p < 0.001 for EC, p = 0.021 for EOF, p < 0.001 for ECF).

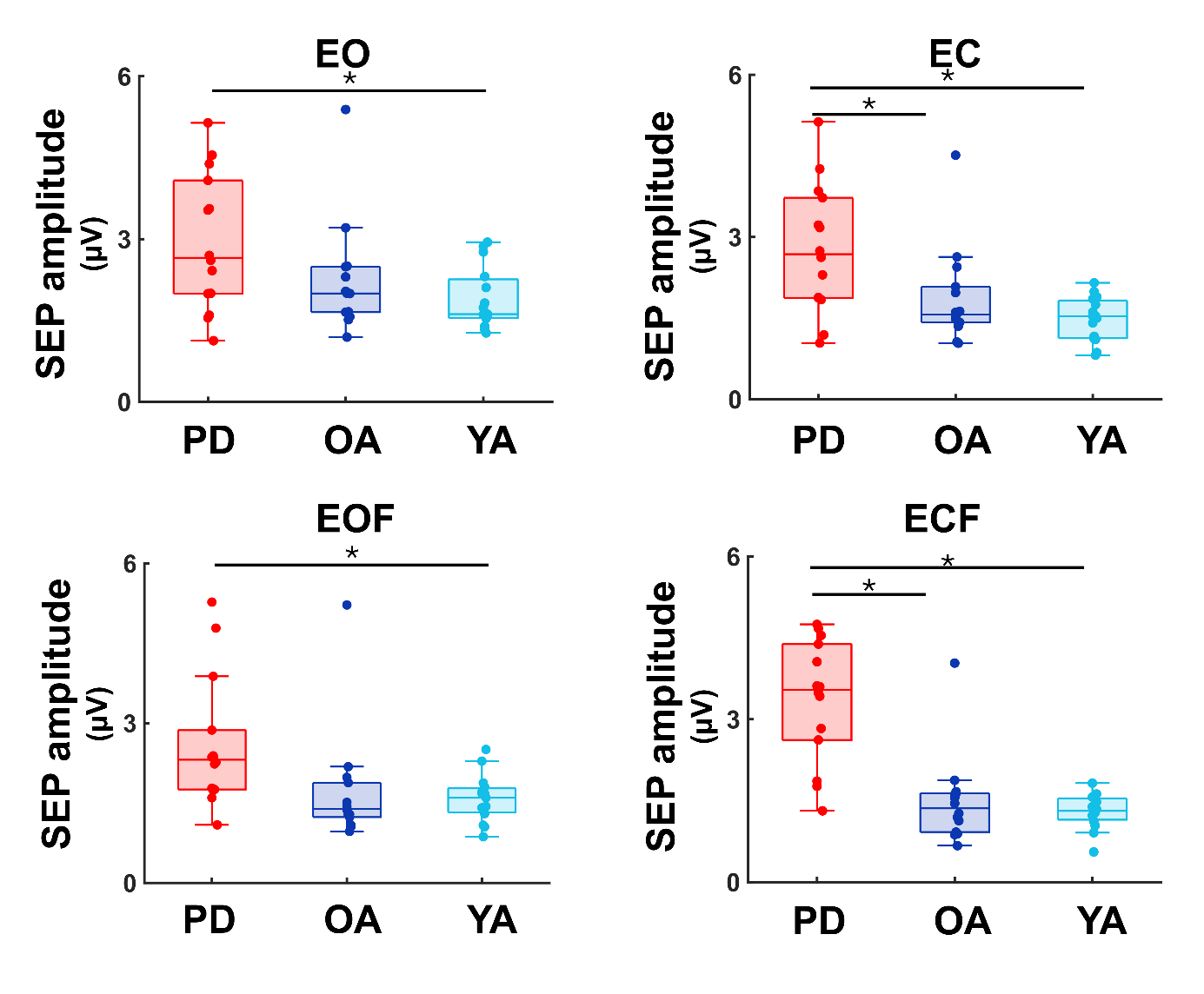

Supplementary Figure 1: Box and whisker plots, with scattered dots indicating each subject showing differences in the SEP amplitude between the three groups, PD (red), OA (dark blue) and YA (light blue) groups across the four balance conditions, eyes open (EO), eyes closed (EC), eyes open on foam (EOF) and eyes closed on foam (ECF) shown in each panel.

Supplementary Figure 2 shows SEP amplitude results across conditions for the three groups. While the results for the PD and YA group are unchanged due to the inclusion of the OA outlier, including the it resulted in the SEP amplitude for EO significantly exceeding EC (p= 0.045), in addition to the EOF and ECF conditions (p < 0.001) that is reported in the main manuscript.

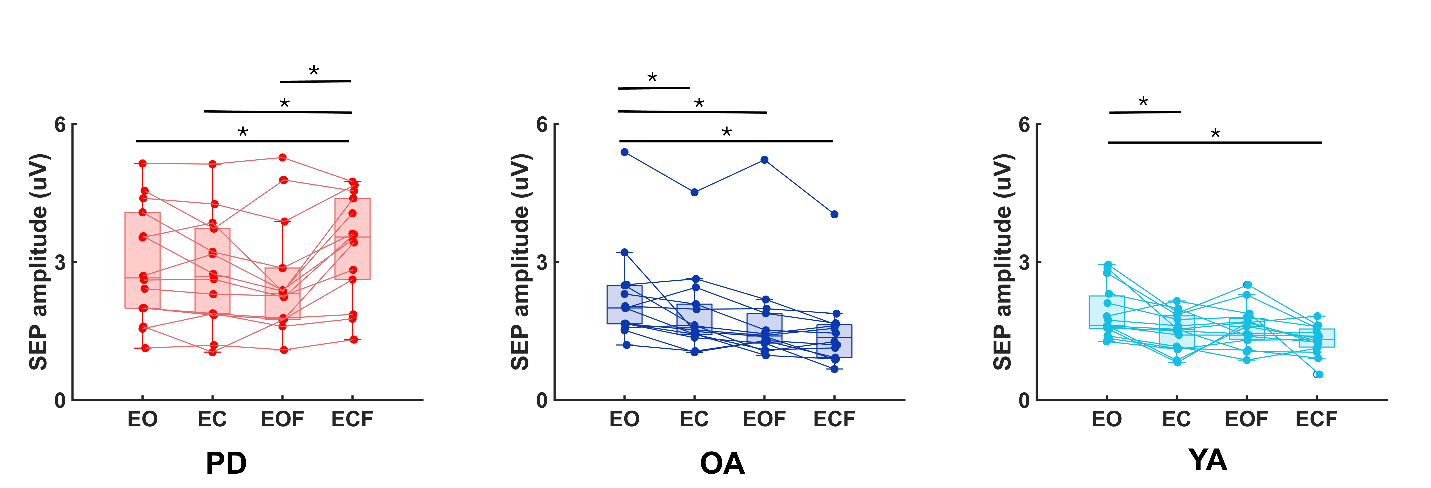

Supplementary Figure 2: Box and whisker plots, with scattered dots indicating each subject showing the trend in the SEP amplitude as the balance conditions become increasingly challenging from eyes open (EO), eyes closed (EC), eyes open on foam (EOF) to eyes closed on foam (ECF) for the three group, PD (red), OA (dark blue) and YA (light blue).
