## Supplementary File 2 for "Impaired Sensory Gating During Standing Balance in Parkinson’s Disease"

We investigated the following time-domain and frequency domain measures for the Center of Pressure (COP).

1. Mean Center of Pressure velocity (COPv), defined as the average speed at which the center of pressure (COP) moves during quiet standing and was calculated as the total distance the COP traveled by the total time.
2. RMS COP A-P and M-L: The root mean square of the COP (RMS COP) was defined as the square root of the arithmetic mean of the squared radius in the A-P and M-L directions.
3. Frequency dispersion (freqdisp): defined as the variability in the frequency content of the power spectrum.

Prior work has found these variables to be associated with aging and with the risk of falling (27-31).

COPv: Results from the 3 (group) x 4 (condition) ANOVA revealed a significant group by condition interaction for 95%COP area (F= 4.671, p < 0.001, ***η***^2^ = 0.172). Pairwise post hoc comparisons using Bonferroni corrections showed that for PD and OA swayed significantly more than YA for EO (p = 0.023 and p = 0.003, respectively), and EC conditions (p < 0.001 and p < 0.001, respectively). While OA did not sway significantly more than YA for EOF (p = 0.139) and for ECF (p = 0.737), PD group showed significantly greater sway than YA for EOF (p = 0.051) and for ECF (p = 0.009). There were no significant differences between the OA and PD group for the EO, EC, EOF, and ECF (p>0.05).

RMS COP M-L: Results from the 3 (group) x 4 (condition) ANOVA revealed a significant group by condition interaction RMS COP M-L (F= 6.381, p < 0.001, ***η***^2^ = 0.221). Pairwise post hoc comparisons using Bonferroni corrections showed that PD and OA swayed significantly more than YA for EO (p < 0.001 and p < 0.001, respectively), and EC conditions (p = 0.006 and p = 0.004, respectively). Both OA and PD did not sway significantly more than YA for EOF (p ~ 1.000 and p= 0.121, respectively), and ECF (p ~ 1.000 and p= 0.182, respectively). There were no significant differences between the OA and PD group for the EO, EC, EOF, and ECF (p>0.05).

RMS COP A-P: Results from the 3 (group) x 4 (condition) ANOVA revealed a significant group by condition interaction RMS COP M-L (F= 2.765, p = 0.029 0.001, ***η***^2^ = 0.109). Pairwise post hoc comparisons using Bonferroni corrections showed that PD and OA swayed significantly more than YA for EO (p = 0.005 and p = 0.005, respectively). For EC, OA did not sway significantly more than YA for EOF (p = 0.130), PD group showed significantly greater sway than YA (p = 0.018, respectively). Both OA and PD did not sway significantly more than YA for EOF (p = 0.325 and p= 0.186, respectively), and ECF (p ~ 1.000 and p= 0.912, respectively). There were no significant differences between the OA and PD group for the EO, EC, EOF, and ECF (p>0.05).

Freqdisp: Results from the 3 (group) x 4 (condition) ANOVA revealed a significant condition effect (F= 4.240, p = 0.010, ***η***^2^ = 0.086) and for group (F= 3.748, p = 0.031, ***η***^2^ = 0.143) but a non-significant group by condition interaction for freqdisp (F= 0.923, p = 0.471, ***η***^2^ = 0.039). Pairwise post hoc comparisons using Bonferroni corrections showed that YA group had significantly greater freqdisp than OA group (p = 0.038) but there were no significant differences between YA and PD (p ~ 1.000) and OA and PD (p = 0.138). Furthermore, freqdisp was significantly greater at ECF compared to EOF (p = 0.023) but there were no differences between other conditions (p>0.05).
